# A pressure sensing protein kinase

**DOI:** 10.1101/435008

**Authors:** Radha Akella, Kamil Sekulski, John M. Pleinis, Joanna Liwocha, Jenny Jiou, Haixia He, John M. Humphreys, Jeffrey N. Schellinger, Jianrui Hu, Melanie H. Cobb, Lukasz Joachimiak, Aylin R. Rodan, Elizabeth J. Goldsmith

## Abstract

Cells respond to hydrostatic pressure to maintain cellular, organ, and organism level functions, yet the direct pressure sensors are largely unknown. Here we show that hydrostatic pressure directly activates <u>W</u>ith <u>N</u>o Lysine(<u>K</u>) kinase-3 (WNK3) ^1^, a soluble intracellular protein kinase. Using gel filtration we demonstrate that pressure induces a dimer to monomer transition in a construct of the <u>u</u>nphosphorylated kinase domain of WNK3 (uWNK3-KDm or uWNK3). The uWNK3 has not been crystallized, but crosslinking data suggest that the uWNK3 dimer corresponds to crystallographically observed dimer of WNK1 (uWNK1-KDm, or uWNK1) ^2,3^. Sequence alignments with WNKs from species living in different pressure environments and mutational analysis lend further support for this idea. Unique features of the uWNK1 structure suggest a mechanism involving bound water. We further show that hydrostatic pressure activates full-length WNK3 in *D. melanogaster* tubules.

Three decades of research on Na^+^Cl^-^ cotransporters (NCCs) and Na^+^K^+^2Cl^-^ cotransporters (NKCCs) suggested the existence of an osmotically activated and chloride inhibited protein kinase to explain NCC and NKCC regulation^4-6^. WNKs are in the pathway for the regulation of NCCs, since anti-hypertensive diuretics that target NCCs are effective in treating familial hypertension associated with WNK isoforms ^7^. WNKs are activated by osmotic pressure in cells in culture, becoming phosphorylated ^8,9^. WNK’s autophosphorylate in low chloride ^3^, as noted above, and thus WNK kinases may be the anticipated osmotically activated and chloride inhibited kinase involved in cotrasporter regulation^10,11^. WNKs are associated with familial forms of both hypertension and hypotension^7,12,13^, giving rise to the possibility that hydrostatic pressure also is a regulator of WNK kinases. We tested the pressure sensitivity of WNK3 because WNK3 knockout reduces intracranial pressure in a stroke model^14^. We observed pressure-induced autophosphorylation of uWNK3. The pressure sensitivity of other WNK isoforms as well as the osmotic pressure sensitivity of WNK3 and WNK isoforms will be addressed in future studies.

WNK isoforms autophosphorylate on a primary activating phosphorylation site in the activation loop {Thastrup, 2012 #4490;Piala, 2014 #4897;Xu, 2002 #4603. WNK3-KDm as expressed in bacteria is fully phosphorylated on this serine, Ser308, as well as on Ser304. Mass spectrometric analysis reveals two significant sites in the S-Tag used for expression (Extended Data Table 1). Dephosphorylation with phosphatases to make uWNK3 removes most of this phosphorylation (Extended Data Table 1).

Hydrostatic pressure induces autophosphorylation of uWNK3 (Fig. 1A). Pressure of 190 kPa (28 psi) was applied with N_2_ gas in an Amicon concentrator, and autophosphorylation was visualized by phospho-staining (Fig. 1A). The pressurized uWNK3 autophoshorylates more than unpressurized uWNK3 on the major sites observed by mass spectrometry (Extended Data Table 1). The maximal enhancement of autophosphorylation of uWNK3 occurred at 7-15 minutes (Fig. 1A). At a 15 min time point, pressure induced approximately 30% increase in phosphorylation of uWNK3 (p<0.01) (Fig. 1B and Extended Data Fig. 1). Autophosphorylation increased enzyme activity. uWNK3 was used to phosphorylate a peptide containing the WNK substrate OSR1 (oxidative stress-responsive-1) recognition site{Anselmo, 2006 #3558;Vitari, 2006 #4527}. Hydrostatic pressure increased the phosphorylation of the GST-fused OSR1 peptide by about 30%, (p<0.0001) (Fig. 1B, Extended Data Fig. 1B). Chloride is inhibitory of WNK1^3^. The chloride opposition to the pressure-induced autophosphorylation of uWNK3 was measured by increasing the [Cl^-^] from 50 mM to 250 mM. The uWNK3 was inhibited more than 50% (p<0.001 and p<0.0001) (Fig. 1C).

**Fig 1.**
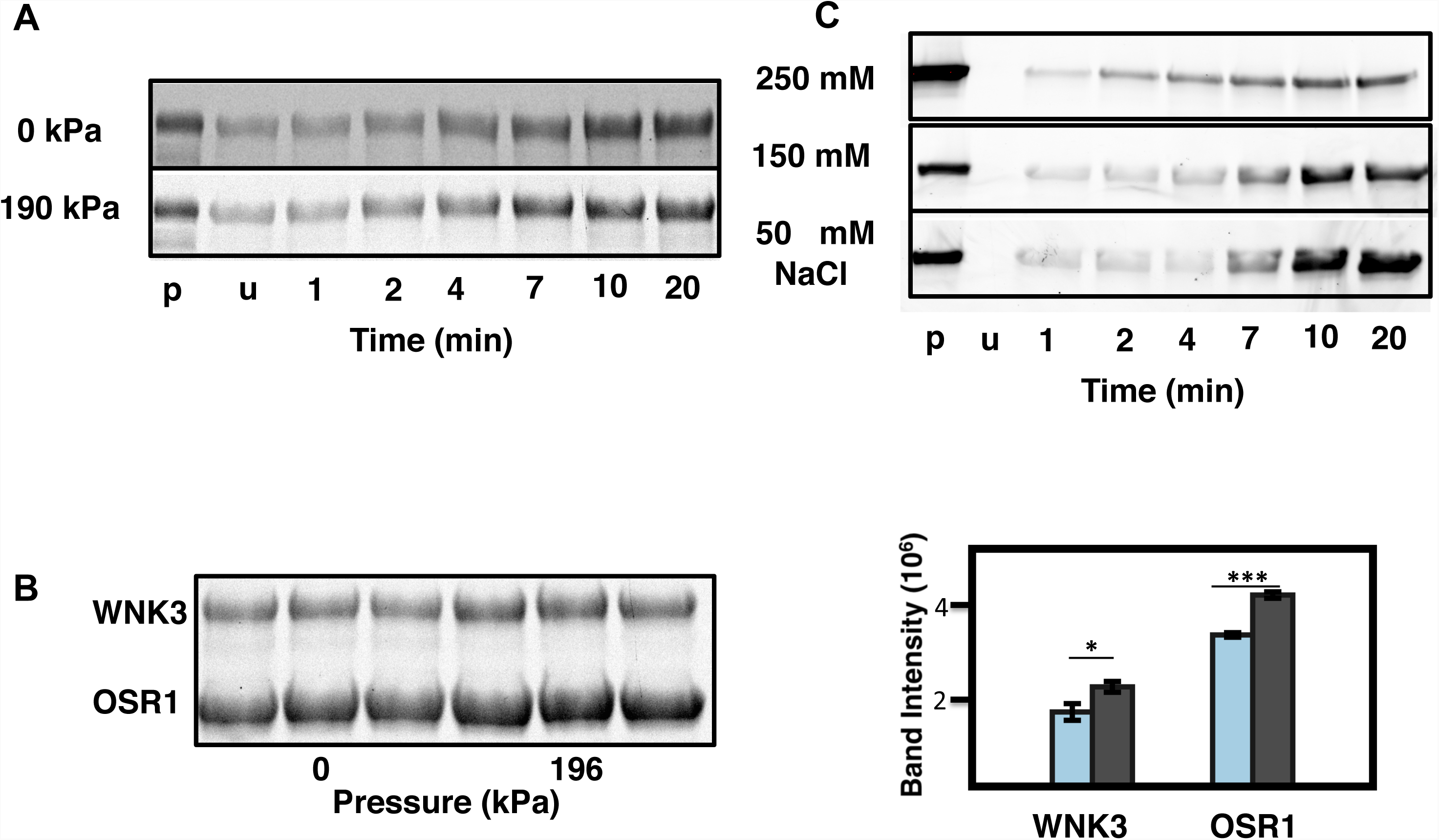
Autophosphorylation of WNK3-KDm, and OSR1 as a function of hydrostatic pressure and chloride. (**A**) Timecourse of autophosphorylation WNK3-KDm with 0 and 190 kPa applied hydrostatic pressure (N_2_) for the times indicated. 4 mM WNK3-KDm, standard conditions, in 150 mM [Cl^-^], 25 °C. Pro-Q stain of phospho-protein. (**B**) Effect of pressure on both WNK3-KDm and GST-OSR1(314-344), 3 replicates each at 0 and 190 psi, 15 min reactions. Light blue, unpressurized WNK3-KDm and GST-OSR1(314-344) phosphorylation, (difference p<.01), gray, pept-OSR1 phosphorylation (p <.0001). (**C**) Chloride opposition of autophosphorylation of WNK3-KDm at 190 kPa at 250 mM NaCl, 150 mM NaCl 50 mM NaCl, at times indicated. One of three replicate gels is shown.

Chloride binds to and stabilizes an inactive, phosphorylation incompetent dimer of WNK1^3^. Gel filtration of uWNK3 in 150 mM NaCl reveals this dimer (Fig. 2A) as well as a higher order aggregate and a monomer. Influence of NaCl on the oligomer state was observed by gel filtration at a lower salt concentration, revealing a shift to a monomer (Fig. 2B). To test whether pressure also induces monomer formation, we carried out gel filtration at a higher back pressure, 1.1 MPa, maintaining the 150 mM NaCl, and again observed a dimer to monomer shift (Fig. 2C). The gel filtration data fit a model of a conformational equilibrium between two states, an inactive dimer and a phosphorylation competent monomer (Fig. 2D). The dimer is promoted by chloride, and the monomer is promoted by pressure.

**Fig 2.**
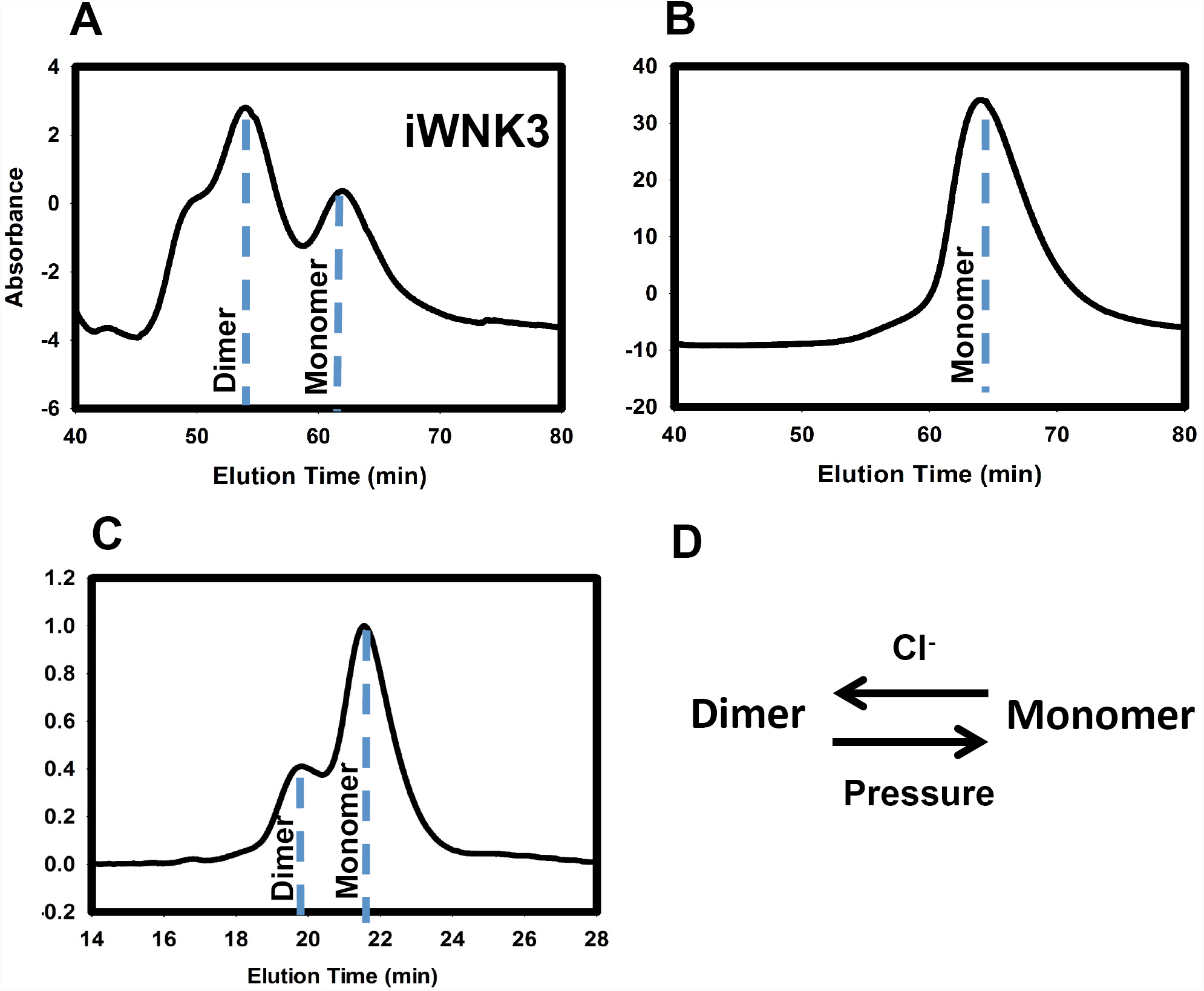
Oligomerization state of iWNK3 as a function of hydrostatic pressure and chloride. (**A**) Size exclusion chromatography of dephosphorylated WNK3-KDm on a Superdex 75 column at a back pressure of 400 kPa, 5mg/ml, (**A**) 150 mM NaCl, (**B**) 50 mM NaCl. (**C**) Size exclusion chromatography at 1MPa back pressure together with with multi-angle light scattering. (**D**) Schematic of dimer to monomer transition, chloride promoting the dimer, pressure the monomer.

In addition to uWNK3, uWNK1 autophosphorylation was tested and found to be slightly pressure sensitive (Fig. 3A, Extended Data Fig. 2). To determine whether pressure-induced auotphosphorylation is unique to WNKs, we tested for pressure sensitivity in the autophosphorylating MAP3Ks TAK1-TAB (TGFβ activated kinase-TAK binding protein 1)^15^ and ASK1 (apoptosis signaling kinase 1) ^16^. The MAP kinase p38 also weakly autophosphorylates. TAK1-TAB autophosphorylates significantly under the conditions used, but is not influenced by pressure (Fig. 3A). The autophosphorylation of ASK1 and p38 were less, but both showed small positive changes with pressure. The action of pressure on the trans-phosphorylation reactions of the MAP2K MEK6 (active mutant) and the MAP2K MEK1 (active mutant) was inhibitory (Fig. 2D, gels in Extended Data Fig. 2).

We also tested whether the phosphatases, PP1cγ, MAP kinase phosphatase-3 (MKP3), λ-phosphatase, and shrimp alkaline phosphatase also respond to pressure. These enzymes showed no pressure-induced activity, as measured by the loss of phosphate from p-nitrophenyl phosphate (Extended Data Fig. 3).

**Fig 3.**
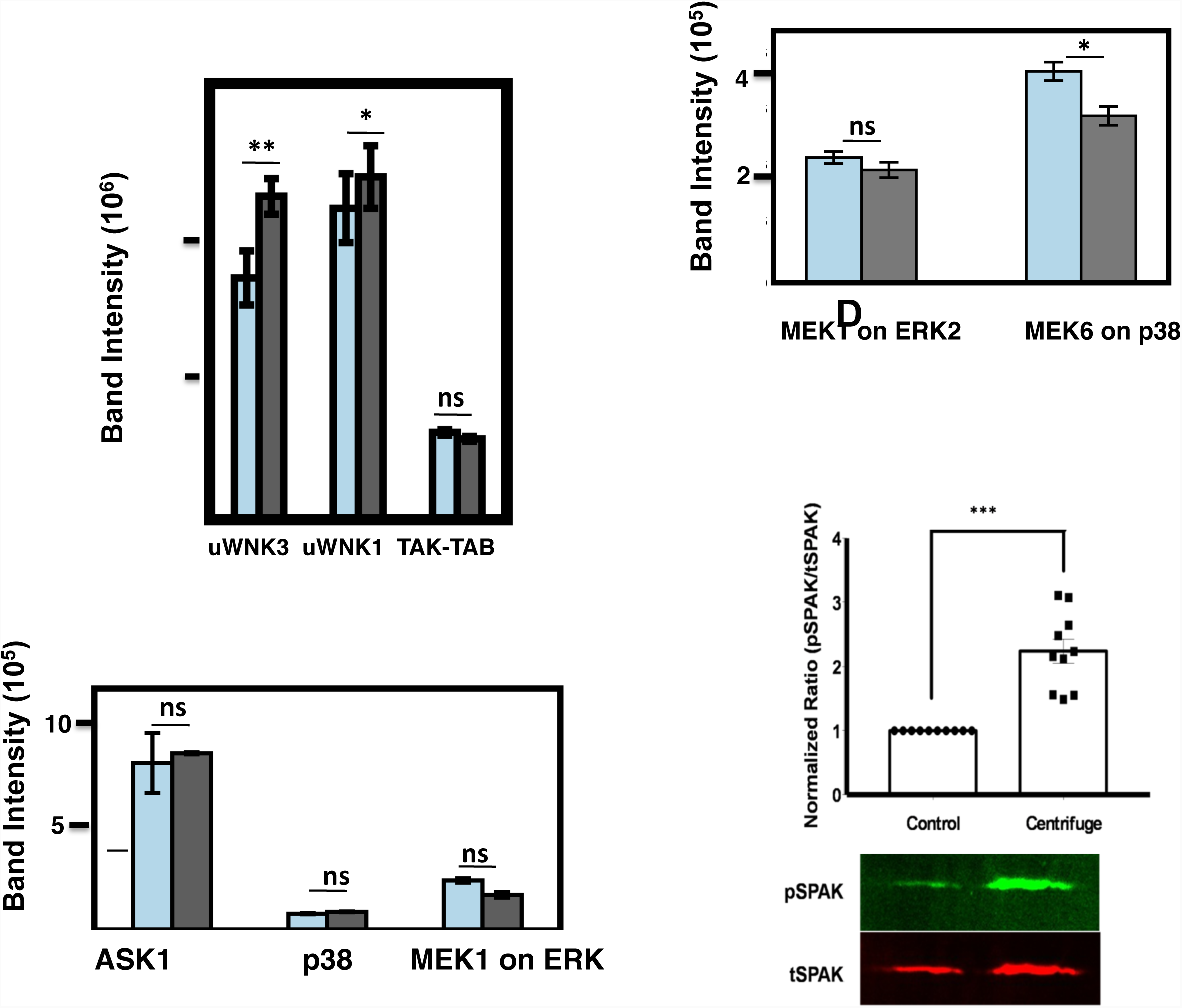
Pressure effects on control kinases, and pressure induced phosphorylation of WNK3 *in vivo*. (**A**) Autophosphorylation of iWNK1 (from Fig. 1B), and TAK1-TAB. (**B)** Autophosphorylation of ASK1 and the MAPK p38, and trans-phosphorylation of ERK2 by MEK1 4mM kinase, 7 min reactions, 150mM [Cl^-^], 25 °C (**C)** Trans-phosphorylation of the MAPK p38 by constitutively active MEK6/DD, with that of ERK2 by MEK1/F53L (active mutant) shown for comparison. Light blue bars in A-C are unpressurized, gray bars, 190 kPa pressure. (**D)** WNK3 pressure activation in *D. melanogaster* renal tubules was expressed in *D. melanogaster* renal tubules in which endogenous *Drosophila WNK* was knocked down and full-length human WNK3 was expressed, together with kinase-dead rat SPAK^D^219^A^ as a WNK3 substrate. Pressure was applied to isolated tubules by centrifugation. SPAK phosphorylation was quantified by Western blot using anti-pSPAK and anti-total-SPAK antibodies. 10 independent experiments, with 30 tubules/condition, were performed. In each experiment, the p-SPAK/t-SPAK ratio in centrifuged tubules was normalized to control to account for day to day experimental variation. ***, p<0.001, one-sample t-test to theoretical mean of 1. A sample Western blot is shown.

We examined the effects of pressure on the activity of WNK3 in *D. melanogaster*. We have previously shown that WNK-SPAK/OSR1 signaling is involved in ion transport in the *Drosophila* Malpighian tubule^17,18^. Full-length WNK3 was expressed together with its substrate SPAK (Ste20/SPS1-related proline/alanine-rich kinase, kinase dead). Isolated tubules were subjected to hydrostatic pressure by centrifugation (experimental scheme shown in Extended Data Fig 4). The activation of WNK3 was measured by increased phosphorylation of the transgenicly-expressed kinase-dead SPAK, under hydrostatic pressure of 80 kPa, lower than that used *in vitro* experiments (Fig. 1). The centrifugation resulted in a 2-fold increase in the ratio of pSPAK (phosphorylated) to tSPAK (total) (p<0.0001).

**Fig 4.**
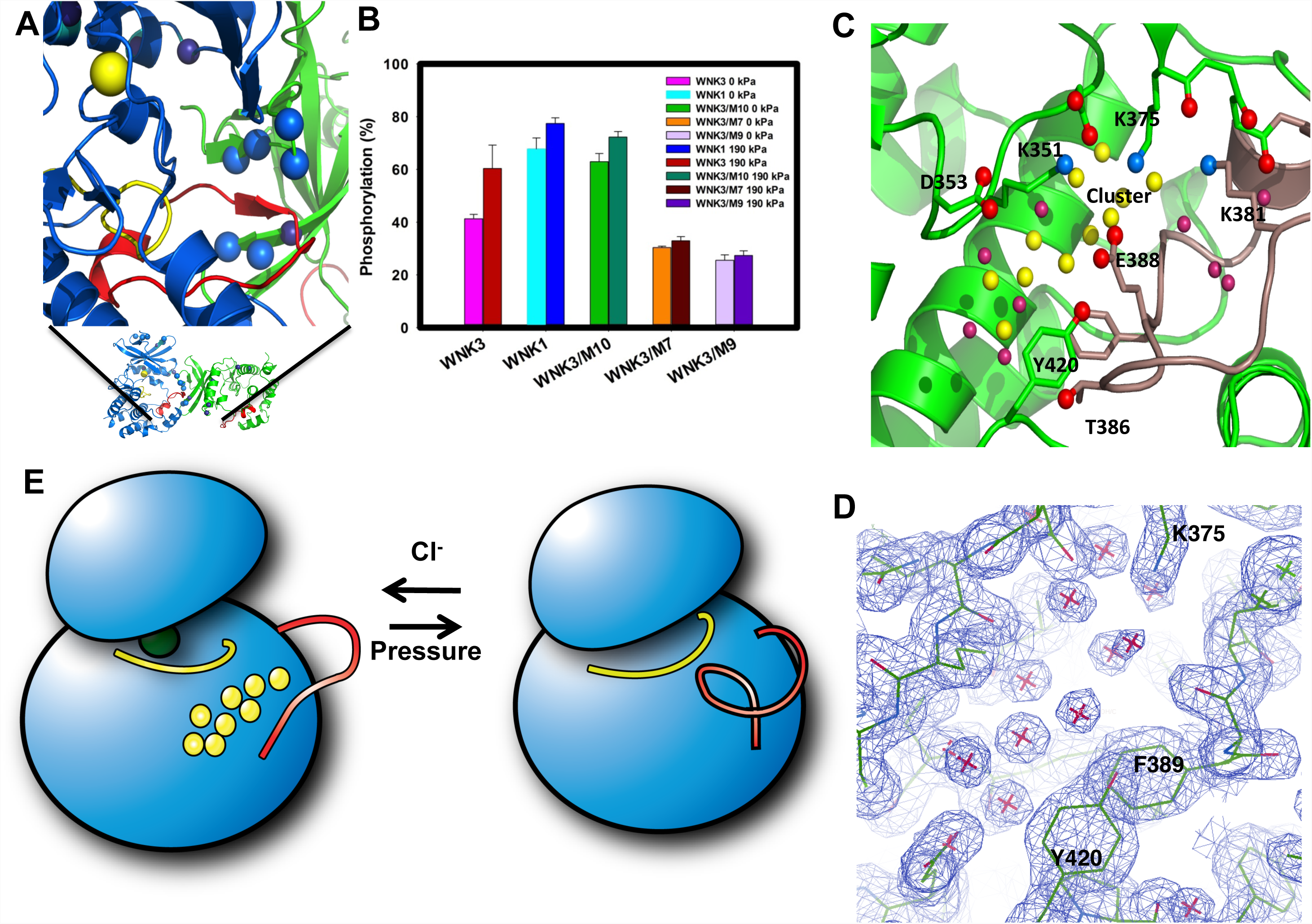
Model and molecular structure of iWNK1/SA. (**A**) Dimer interface in the crystal structure of uWNK1A. Marine spheres indicated positions mutated, dark spheres represent other positions different between WNK3 and deep sea fish WNK1 (*N. coriiceps)*. (**B**) Autophosphorylation of uWNK3, uWNK1, and mutants, without and with 190 kPa added pressure. The mutants are M10 (A139G, I216P and L217S) M9 (A139G, T140M, P142N, I216T and L217V) and M7 (I216P, L217S) (**C**) Cluster of charges in Subunit A that form a large cavity Water molecules underneath the charges is yellow. (**D**) Electron density for the water molecules 3FPQ contoured in COOT at 1 s. (**E**) Model for conformational changes in one subunit based the structure uWNK1A, 3FPQ, and pWNK1, 4PWN, showing water molecules lost as the activation loop refolds.

One structure of a dimeric inactive WNK is available, uWNK1 (uWNK1(Ser382->A), uWNK1A) ^2^. uWNK3 is expected to have a similar structure as uWNK1A based on the >90% sequence identity. Sequence alignments with selected WNKs, we hypothesize might have altered pressure sensitivity due to a difference in habitat pressure, revealed that amino acid replacements mapped of the dimer interface of uWNK1A (Fig. 4A).

Mutational analysis was used to test the dimer in pressure sensing. Mutations in WNK3 were introduced to to resemble the interface of WNK1 or that of WNK1 from *N. coriiceps* (fish) (mutants made are described in Methods, results in Fig. 4B, sequence alignment in Extended Data Fig. 5A). Of the three mutants analyzed, one was more active and two were less active, and all three had reduced pressure sensitivity (Fig. 4B). The lack of activity of mutants M7 and M9 may be due to changes in stability (data not shown). Further evidence for the uWNK1 dimer in uWNK3 was obtained from lysine crosslinking with disuccinylsuberate (DSS) in the absence and presence of 190 kPa pressure. Crosslinks were analyzed by mass spectrometry (See Methods) (Extended Data Tables 2 and 3). A distinct constellation of crosslinks was observed under the two conditions. In the absence of pressure, some of the crosslinks specific to the unpressurized form clearly arise from the dimer (Extended Data Table 2A, see Extended Data Fig. 5B for how crosslinking was assigned to monomer or dimer).

It is intriguing to consider how uWNK3 might sense pressure. Dimeric uWNK1A is unusual in having extensively remodeled activation loops in both subunits (from the ^368^DLG sequence through the ^386^APE sequence in human WNK1) (Fig. 4A), giving rise to a large cavity at the active site (Extended Data Fig. 6A). The cavity volume calculated in POCASA ^19^ gave a volume of 1200Å^3^ for uWNK1A as compared with the cavities in a typical active protein kinase such as PAK6 (p21 activated kinase-6, 450Å^3^) (Extended Data Fig. 6B). The activation loop and catalytic loop interact with each other forming an ionic cluster (Fig. 4C) rather than the canonical hydrophobic interactions observed in typical protein kinases (Extended Data Fig. 6C) ^20,21^. Two residues in the cluster, D349 and K351 (WNK1 numbers) are in the catalytic loop and pan-kinase conserved, others are pan-WNK conserved, D353 in the catalytic loop and K375, and K381 and E388 in the activation loop (Fig 4D). Well-ordered water (Fig. 4C,D) is trapped beneath the ionic cluster (Fig. 4D). Similar ion pairs and waters are present in other inactive WNK1 structures, such as 5DRB ^22^. A model for what might happen to one subunit on going into an active configuration is shown in Fig. 4E, and is based on structure of pWNK1 and other active kinases. Extra ion pairs occur throughout the structure (not shown), and is reflected in the composition of WNK1 and WNK3, which is 10% lysine and 10% glutamic acid, twice as proteins of comparable size ^23^. Thus, we think that this cluster of charges and other ion pairs may be trapping water.

The data presented here demonstrate that a soluble protein kinase is directly activated by hydrostatic pressure *in vitro* and *in vivo*. The enhancement of activity is modest, about 30%, but this increase is amplified by the cascade, phosphorylation of the downstream substrate. The pressure applied was well above the 16 kPa (120 mmHg) typical of systolic blood pressure, but orders of magnitude below pressures often used to study effects on proteins (200 MPa and more)^24-26^. The pressure sensing is an intrinsic property of the kinase domain and involves a conformational equilibrium between an inactive dimer and an autophosphorylation-competent monomer. We hypothesize a role for bound water in pressure sensing based on the structure. The apparent involvement of water is reasonable because water on the surface of proteins is lower density than bulk water ^27-30^. Thus, applying pressure should disfavor the configuration with the most bound water.

The direct pressure sensing of uWNK3 has not been anticipated. WNK3 is involved in volume regulation and neuronal excitability in brain ^14,31-33^. Further experimentation is required to determine if WNK3 pressure sensing is involved in these processes. Other WNK kinases may be sensitive to pressure also, and should be characterized the context of full length proteins. The apparent regulation of WNK kinases by hydrostatic pressure may account for familial hypertension and hypotension associated with WNKs ^13^. The present study expands the field of pressure sensors from membrane bound proteins to include molecules in the cytoplasm.

## Acknowledgements

We thank the American Heart Association (14GRNT20500035 (EJG) and 16CSA28530002 (EJG and ARR)), NIH DK110358 (ARR and EJG) and the Welch Foundation (I1128, EJG and I1243, MHC) for support. We thank Steven McKnight for discussion of this work, Kim Orth and Kelly Servage for help with mass spectrometry of crosslinking data collection. We thank Tung-Chung Mou, University of Montana, for SEC/MALS data collection and analysis.

## Author’s Contributions

R.A. made the initial discovery, planned experiments, performed data analysis and re-refinement of structure, and assisted with manuscript preparation. K.S. collected most of gel-based data presented. H.H. performed all DNA work. J.L. performed early characterization of pressure effects. J.J. designed pressure sensitivity mutants based on sequence alignments. J.M.H. performed mass spectrometry. J.M.P., J.N.S and J.H. conducted pressure experiments in *Drosophila* Malpighian tubules. M.H.C. provided OSR1 plasmid, and assisted with the manuscript. L.J. assisted crosslinking data analysis. A.R. R. planned and oversaw tubule experiments, and assisted with manuscript preparation. E.J.G. initiated and guided this work and wrote the manuscript.

---

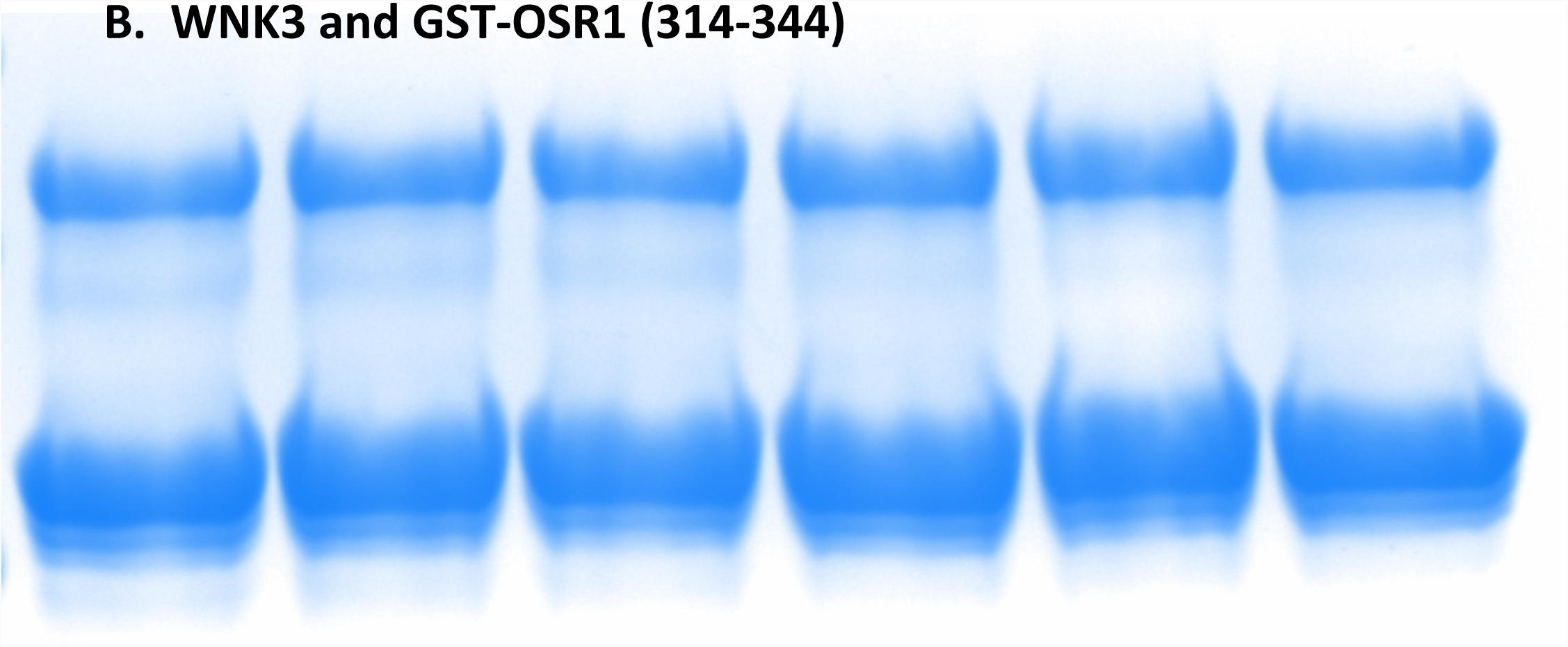

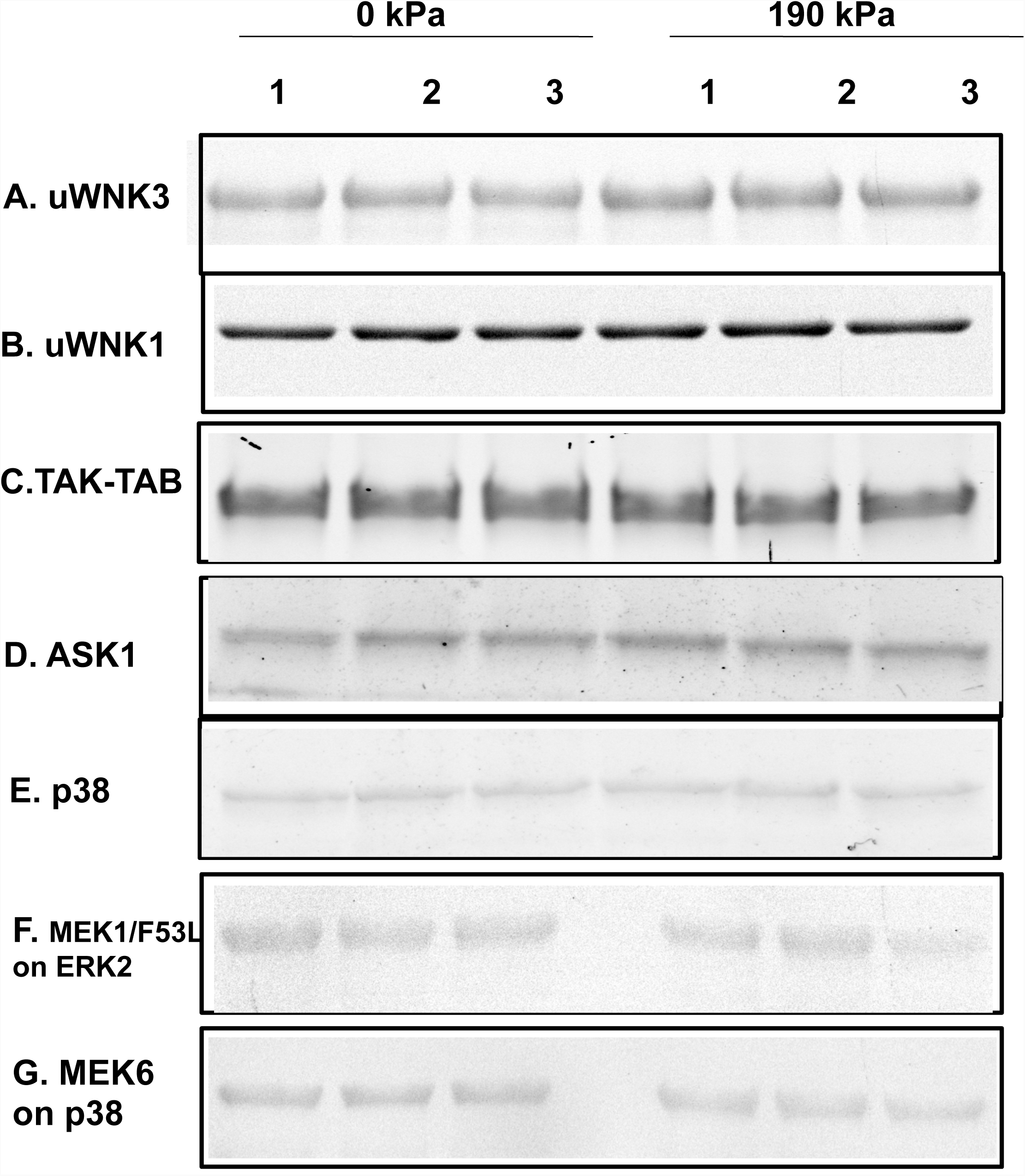

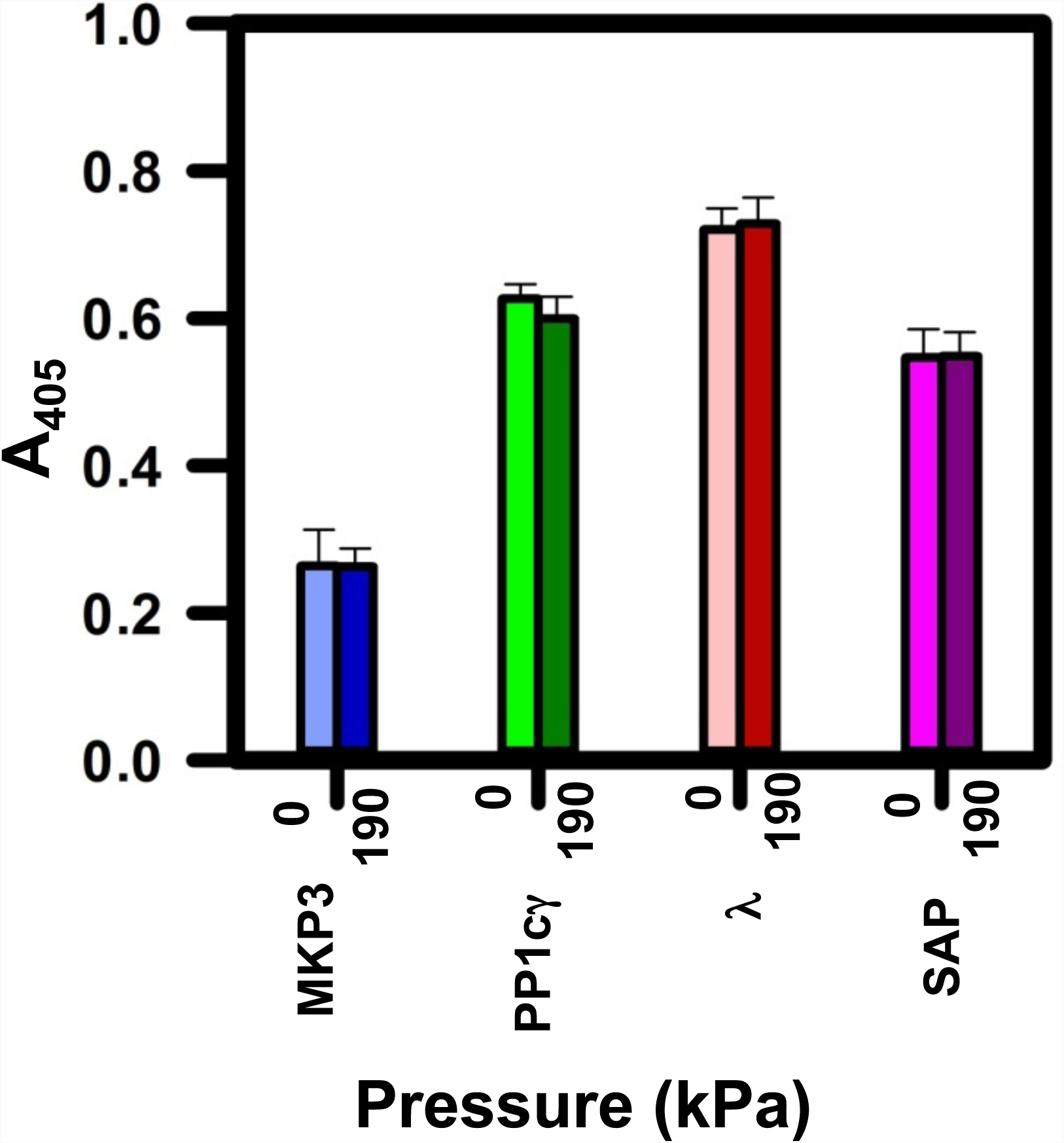

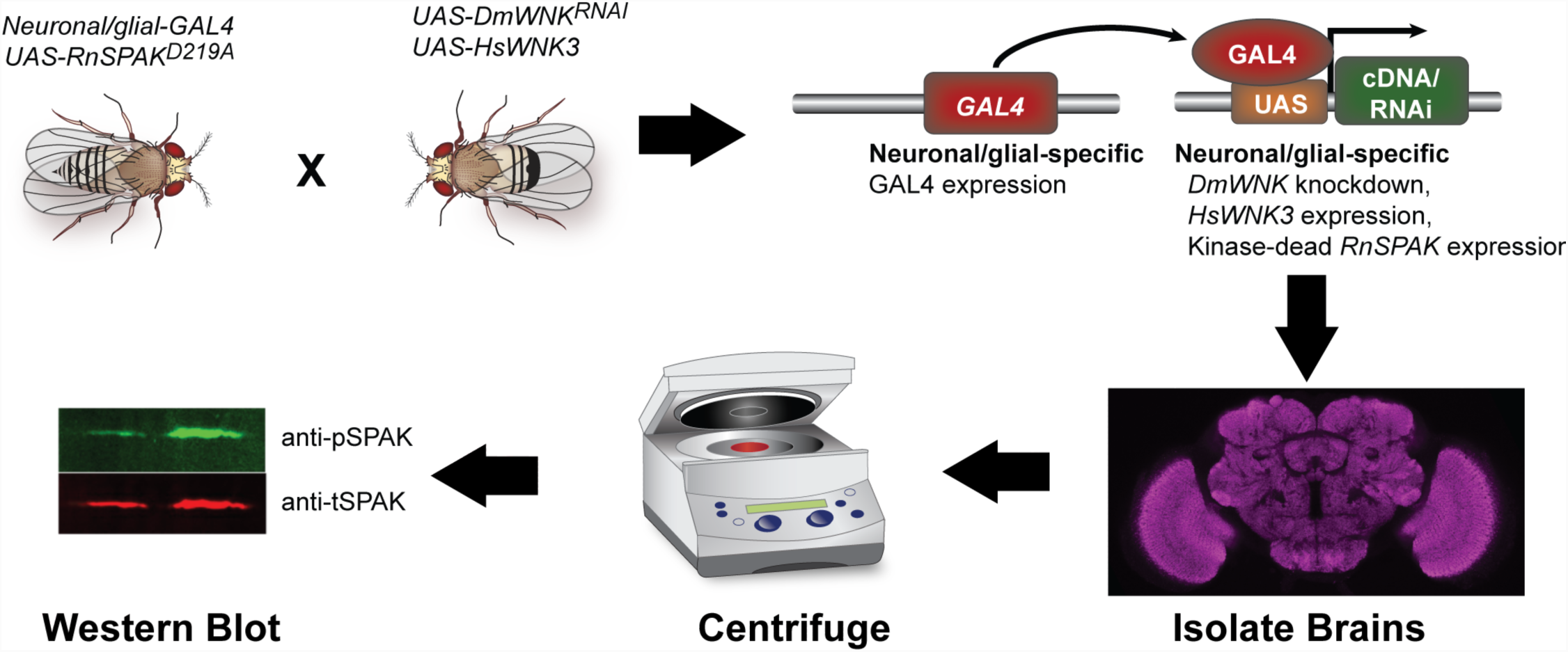

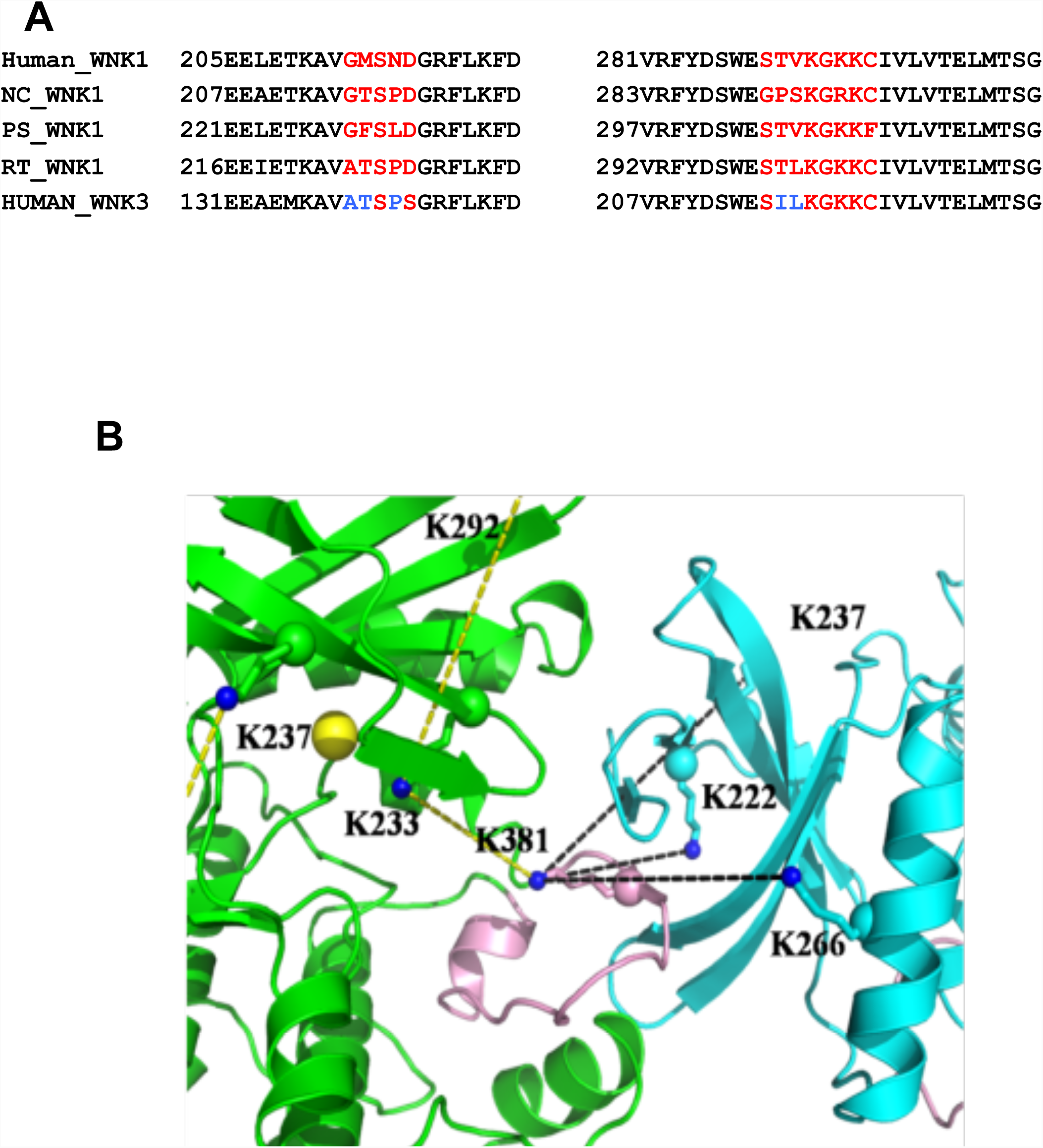

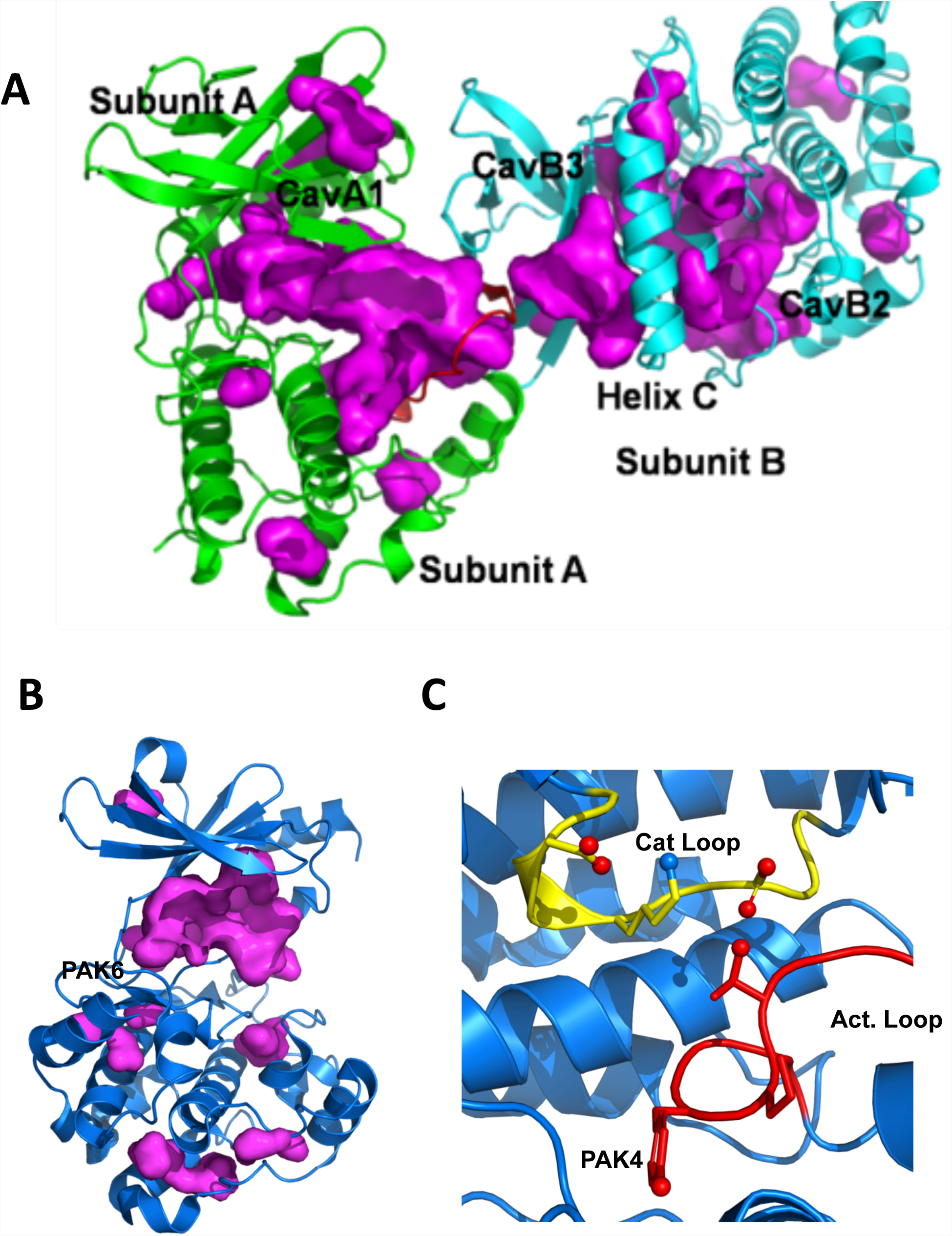

