## Supplemental Methods and Tables for "A pressure sensing protein kinase"

**This file includes:**
Methods
Legends Extended Data Figs. 1-6 (Figs in with main text)
Extended Data Tables 1-3
References

### Methods

##### Protein reagents

The sequence for human WNK3-KDm (118-409) was optimized to eliminate rare codons by Genscript Inc. (New Jersey) and subcloned into a pET29b vector. The clone for WNK1-KDm(194-483) was described previously [1]. A vector for ASK1 was obtained from Stephan Knapp, Oxford structural Genomics Consortium. A vector for PP1cγ was a gift of Anna Depaoli-Roach[2]. Expression of proteins was conducted similarly in *E. coli* using Terrific Broth (Fisher

Scientific), induced with 0.5 mM isopropyl-thiogalactopyranoside (IPTG) at O.D. 0.8-1, and grown for 18 hours at 18°C. Purifications used Ni-NTA affinity chromatography followed by anion exchange on Mono-Q (10/10) (GE Healthcare); details available in[1,3,4]. The proteins p38, MEK6, and OSR1 were unphosphorylated as expressed in *E. coli*. Unphosphorylated TAK-TAB, expressed in insect cells, was obtained purified from Kenneth Westover[5]. WNK3 and ASK1 were phosphorylated as expressed and were dephosphorylated using phosphatases. WNK3 was dephosphorylated using PP1c$\gamma$ at a10:1 ratio. ASK1 were dephosphorylated using $\lambda$–phosphatase (Santa Cruz Biotechnologies, Inc.) and shrimp alkaline phosphatase (Invitrogen) in 10:1 and 5:1 molar ratios respectively.. All phosphatase reactions were conducted in 0.5 mM $MnCl_2$, at 4°C overnight. Phosphatases in the above reactions were removed using Ni-NTA, based on the His-tags on WNK3-KDm and ASK1. Kinases was buffer exchanged into 50 mM Hepes, pH 8.0, 150 mM NaCl, and 1 mM dithiothreitol[6]. Bacterial expression of GST-tagged Human OSR1 (314-344) was similar, inducing with 0.4 mM IPTG at O.D. 0.6. The GST-OSR1(314-344) was purified on a glutathione sepharose column (GE Healthcare) and buffer-exchanged into 20mM HEPES, pH 7.4, 50 mM NaCl and 5% glycerol.

**Autophosphorylation Assays**

Autophosphorylation assays were carried out in 20 mM HEPES, pH 7.4, 20 mM $MgCl_2$, 2.5 mM ATP at 25 $^{o}$C. Reactions were initiated by the addition of the 4 $\mu$M or 6 $\mu$M de-phosphorylated enzymes, iWNK3, ASK1-KDm, and TAK-TAB, GST-OSR1 and p38 were used. All reactions were stopped by the addition of guanidine hydrochloride to 1 M or Laemelli SDS gel running buffer. Protein phosphorylation was measured in gels with a stain for phosphorylated amino acids, Pro-Q Diamond (Life Technologies Inc.) or by mass spectrometry.

**Trans-phosphorylation Assays**

Transphosphorylation assays were carried out in 20mM HEPES, pH 7.4, 20 mM $MgCl_2$, 2.5 mM ATP with an enzyme: substrate ratio of 1:100 at room temperature and run for 7 minutes. Reactions were initiated by the addition of ATP. MEK6/DD, constitutively active MEK1 (F53L), and B-Raf phosphorylated their respective substrates GST-OSR1 for both WNK1 and WNK3, p38$\alpha$, ERK2 and MEK1. Reactions were stopped by the addition of Laemelli. Protein phosphorylation was measured using gels stained with Pro-Q Diamond (Life Technologies Inc.).

**Applying pressure and running reaction**

Hydrostatic pressure was applied in an Amicon concentrator connected to a $N_2$ tank fitted with a Boclow low pressure regulator (BOC Corp. UK). Samples were subject to different pressures (added pressure from 0-290 kPa).

**Gel-based phosphostaining**

To detect phospho-proteins, 12% precast BioRad SDS-PAGE gels were run followed by staining with Pro-Q Diamond phospho-protein stain (Life Technologies Inc.), using the manufacturer's protocol. Aliquots for triplicated unpressurized and pressurize reactions were loaded on the same gel. Gels were imaged using a BioRad ChemiDoc MP Imaging System. ProQ Diamond stain signal intensity was observed using the Epi Green protocol and 605/50 filter, and data was quantitated using both Image Lab and ImageJ software[7]. Time course data were collected on the same day, and processed similarly. Time course images reported were scaled by the exposure time.

#### Statistics

Standard deviations from the triplicated Image Lab or ImageJ measurements and t-test p-values comparing unpressurized and pressurized samples were calculated in Excel. * indicates a t-test of <0.1, ** <0.01, and ***<0.0001.

#### Phosphatase reactions

Phosphatase assays were conducted using a phosphatase assay kit (GE Biosciences, St. Louis, Mo.) that utilizes on the difference in extinction coefficient between *p*-nitro phenylphosphate and *p*-nitrotryosine at 405 nm. Assays with and without pressure used 6 μg phosphatase and were run for 30 min at 25°C before being stopped by addition of 3 N NaOH. *p*-nitrotryosine absorbance at 405 nm was measured using a Beckman EU640 spectrophotometer.

#### Mass spectrometry

Proteins were digested in an 8:1 molar ratio with sequence-grade chymotrypsin (Roche) in the presence of 100 mM Tris pH 8.0, and 25 mM $CaCl_2$ (100 $\mu$L total reaction volume) at 30°C overnight. Following digestion, the peptide mixture was separated by HPLC (Agilent 1100) on a RP-C18 column (Phenomenex Aeris Widepore 150 x 2.1 mm) using an acetonitrile-water gradient from 4% to 28% acetonitrile containing 0.1% formic acid throughout. Mass spectrometric analysis was performed on an LCQ DECA XP ion-trap (ThermoFinnigan) with the HPLC coupled inline to an orthogonal electrospray ionization source. MS detector responses were obtained by integration under ion traces corresponding to *m/z* ranges for activation loop peptides. MS/MS spectra were acquired in a data dependent mode and analyzed using MASCOT software (Matrix Science Ltd.)[8,9] or MassMatrix software [10].

The MS/MS fragmentation pattern used to observe pWNK1 phosphorylation was previously reported[11]. pWNK3 gives a similar fragmentation pattern. As purified from bacteria, WNK3-KDm is ~95% phosphorylated on the activation loop sites S304 and S308. Two significantly phosphorylated sites are observed in the S-tag (KET*A T) and thrombin cleavage site (PRGS*M). Four additional  sites showed minor phosphorylation at S125, S289 S363, and in the C-terminal linker to the TEV recognition site.  Dephosphorylation reactions went to completion on S308, and both of the significant tag sites (T4') and( S27').  S304 was left ~30% phosphorylated.  A 15 minute phosphorylation reaction led to rephosphorylation of S308 to ~80%, with 50% phosphorylation on S304, and ~40% on the two sites in the N-terminal tag/thrombin.  In comparison, the catalytically dead WNK3 S208A shows no phosphorylation within the WNK3 sequence, and <1% phosphorylation at the TEV recognition site, as expressed in bacteria.

**Disuccinylsuberate crosslinking**

Disuccinylsuberate (DSS) crosslinks lysine residues[12]. Samples were unpressurized or pressurized at 190 kPa in the Amicon concentrator/$N_2$ setup. Reaction conditions were 4$\mu$M WNK3-KDm, 150 mM NaCl, 10 mM HEPES pH 8.0 and 1 mM DSS in 25 μL for 20 mins at 25°C. Reactions were quenched with 25 mM Tris-HCl pH 7.4. The samples were treated with 6 M guanidine-HCl, to a final concentration of 1M and then proteolyzed with chymotrypsin overnight. Peptides were observed by mass spectrometry on an Orbitrap fusion lumos (ThermoFinnigan). Crosslinks were identified using xQuest/xProphet software[13].

**Gel Filtration and SEC/MALS**

Gel filtration was carried out on a Sephadex-75 column on an AKTA FPLC (GE Healthcare) with a back pressure of 0.3 mPa. SEC/MALS (size-exclusion chromatography followed by multi-angle light scattering) was conducted on a TREO II system, Wyatt –Technology instrument with a back pressure of 800 kPa.

**Differential Scanning Fluorimetry**

Differential scanning fluorimetry (DSF) of WNK3-KDm, WNK1-KDm and mutants were conducted to measure differences in protein stability. 25 μL of 5 μM protein, 50 mM HEPES pH 7.5, and 2.5x Sypro Orange (Life Technologies) of reaction solution was added to a 96 well BioRad Multiplate clear PCR plate. The plate was covered with a BioRad Microseal. Plates were read in a BioRad CFX96 Real-Time PCR system. The temperature was increased from 4 °C to 85 °C, with fluorescence measurements taken every 0.5° C in the 6-FAM fluorescein channel. We determined protein melting temperature ($T_m$) as the inflection point of the negative first differential of the relative fluorescence unit (RFU) with respect to temperature (-d(RFU)/dT).

**Re-refinement of 3FPQ**

Water molecules were added to the model of iWNK1A in COOT[14] based on the |Fo-Fc| and |2Fo-Fc| maps. Restrained refinement of iWNK1A was conducted using REFMAC5 in the CCP4 suite[15] including TLS (Translation-Libration-Screw). The modified structure was deposited (PDB file 6CN9).

**Displaying cavities and calculating volumes**

Cavities displayed in Fig. 4A were calculated in Pymol (Schroedinger, Inc.), using Cavitycull, with a detection radius of 5 solvent molecules. Cavities in the re-refined WNK1-KDm structure were also calculated using Pocket Cavity Search Application (POCASA 1.1, altair.sci.hokudai.ac.jp/g6/service/pocasa)[16] using a probe radius of 2 Å and grid-size of 1.0 Å.

**Generation of transgenic UAS-HsWNK3**

The UAS-HsWNK3 (*Homo sapiens* WNK3) plasmid was generated using the Gateway cloning method (ThermoFisher) with Platinum Pfx DNA polymerase (ThermoFisher 11708013), pENTR/D-TOPO Cloning Kit (ThermoFisher, K240020), LR Clonase II (ThermoFisher, 11791020), and the Gateway compatible destination vector pUASg.attB, obtained from Johannes Bisch and Konrad Basler (Zurich, Switzerland)[17]. Template vector pCMV7.1-hWNK3.2 was obtained from Chou-Long Huang (UT Southwestern, Dallas, TX). HsWNK3.2 was amplified by PCR using primers 5' CACCATGGCCACTGATTCAGGGGATCCAGC 3' and 5' TTTAGGACCAGGAGGGATTGTGGCAGG 3'. The PCR amplicon was gel purified and incorporated into the attL containing entry vector pENTR via directional TOPO cloning per manufacturer's protocol. Full-length pENTR clones were then shuttled into the attR containing destination vector pUASg.attB by LR clonase reaction and sequence confirmed. Midiprep DNA was sent to Rainbow Transgenic Flies for microinjection into stock line #24483 (M{vas-int.Dm}ZH-2A, M{3xP3-RFP.attP}ZH- 51D). Transgenic lines were generated from single male transformants, and PCR confirmation of UAS-HsWNK3 was performed using sequence-specific primers. Transgenic lines were outcrossed for 5 generations to the Rodan laboratory *wBerlin* genetic background.

***In vivo* phosphorylation of SPAK in *D. melanogaster***

Endogenous *Drosophila WNK* was knocked down using *UAS-DmWNK*$^{RNAi}$ (Bloomington *Drosophila* Stock Center, stock #42521 [18], and full-length human *WNK3* was expressed in the principal cell of the *Drosophila* renal tubule, together with the WNK substrate, kinase-dead rat *SPAK*$^{D219A}$, using *c42-GAL4*[19]. The genotype used was *w; UAS-DmWNK*$^{RNAi}$*UAS-HsWNK3/UAS-SPAK*$^{D219A}$*; c42-GAL4/+*.15 pairs of tubules were dissected from 4-6-day-old adult females, transferred to 300 μL of bathing solution for one hour, and then centrifuged at relative centrifugal force (rcf) = 250 for 10 minutes (Eppendorf Centrifuge 5424, 1632 rpm) (estimated pressure, 11.7 PSI), or were subjected to an equivalent temperature gradient in a PTC-100 Programmable Thermal Contoller (MJ Research, Inc). The temperature gradient was determined by centrifuging two separate sets of tubules and measuring temperature at different time points, and increased by a total of 0.75 °C over the first two minutes, beginning at room temperature (~23 °C). Bathing solution consists of a 1:1 mix of saline (in mM, NaCl, 104; $CaCl_2$, 5.5; $MgCl_2$, 20; $NaHCO_3$, 17; $NaH_2PO_4$, 7.5; HEPES, 22; glucose, 35; K gluconate, 40; NMDG gluconate, 125; and Na gluconate, 51.5) and Schneider's medium (in mM, glycine, 3.33; L-arginine, 2.3; L-aspartic acid, 3.01; L-cysteine, 0.496; L-cystine, 0.417; L-glutamic acid, 5.44; L-glutamine, 12.33; L-histidine, 2.58; L-isoleucine, 1.15; L-leucine, 1.15; L-lysine hydrochloride, 9.02; L-methionine, 5.37; L-phenylalanine, 0.909; L-proline, 14.78; L-serine, 2.38; L-threonine, 2.94; L-tryptophan, 0.49; L-tyrosine, 2.76; L-valine, 2.56; β-alanine, 5.62; $CaCl_2$, 5.41; $MgSO_4$, 15.06; KCl, 21.33; $KH_2PO_4$, 3.31; $NaHCO_3$, 4.76; NaCl, 36.21; $Na_2HPO_4$, 4.94; α-ketoglutaric acid, 1.37; D-glucose, 11.11; fumaric acid, 0.862; malic acid, 0.746; succinic acid, 0.847; trehalose, 5.85; and yeastolate, 2000 mg/L) that was then diluted to 300 μL saline/Schneider's mix by adding 277 μL $H_2O$. After centrifugation, tubules were lysed in 2X Laemmli buffer. 50 μL lysate was used to detect phospho-SPAK or total SPAK by Western blotting using 1:1000 dilution of antibodies to phosho-SPAK (Ser373)/phospho-OSR1 (Ser325) (Millipore, Cat. #07-2273, Lot#2840398), total-SPAK (GeneTex anti-STK39 [2E10], Cat.#GTX83543, Lot#821703924) and actin (JLA20-s) (DSHB).These antibodies do not react to antigen in the absence of transgenically expressed SPAK [18]. Secondary antibodies, used at 1:2500 dilution, were fluorophore-conjugated anti-rabbit AzureSpectra 800 (VWR, Cat. #AC2134, Lot #170124-06) or anti-mouse AzureSpectra 700 (VWR, Cat.#AC2129, Lot #170403-07). Protein bands were visualized using a c600 (Azure Biosystems) and quantified in ImageJ by manually outlining the bands and subtracting background pixel intensities from a nearby region. Data were analyzed in GraphPad Prism, version 7.

**Calculation of tubule pressure**

Tubule pressure was calculated using the following equation: Pressure (Pascal) = Force (Newtons) / Surface Area ($m^2$). Force was calculated using the following equation: Force (Newtons or $kg*m/s^2$) = mass (kg) * acceleration ($m/s^2$). The mass of 80 anterior tubules was determined to be 0.0003 g (using AB54-S/FACT Mettler Toledo Balance). Converting this mass to kg per tubule gives $3.75 \times 10^{-9}$ kg/tubule. Acceleration was calculated by the following equation (per centrifuge specifications): Acceleration ($m/s^2$) = relative centrifuge speed (RCF) * g ($m/s^2$) * 1000, where RCF = 250. Thus, the force was determined to be $9.20 \times 10^{-3}$ Newtons. In order to determine the surface area, we assumed tubules to have a cylindrical shape, and that the potential space of the lumen is obliterated under pressure. The outer diameter of the tubule is 35 μm, the inner (luminal) diameter is 17 μm[20], and therefore, under pressure, tubule diameter is 18 μm. Surface area was calculated by the following equation: ($2*\pi*r^2 + 2*\pi*r*h$), where r = radius = 0.000009 m for the tubule, and h = height = 0.002 m for anterior tubules[20]. Thus, the calculated pressure = 80.9 kPa.

**Fig. S1. Coomassie gels of pro-Q diamond stained gels of pressure activation assays.** (**A**) Progress curve (Fig. 1A) (**B**) uWNK3 triplicated unpressurized and pressurized autophosphorylation and phosphorylation of GST-OSR1(314-344) (Fig. 1B). (**C**) Commassie gels of the autophosphorylation of uWNK3 over a time course as opposed by chloride (Fig. 1C).

**Fig. S2. Pro-Q Diamond stained gels of controls.** Pro-Q diamond gel for autophosphorylation reactions (**A**) uWNK3, (**B**) uWNK1, (**C**) TAK-TAB, (**D**) ASK1, (**E**) p38, and transphosphorylation reactions (**F**) MEK1/F53L (active mutant) on ERK2 and (**G**) MEK6/DD (active mutant) on p38 (kinase dead mutant K53R). Autophosphorylation in triplicate. Images adjusted to be visible and not on the same scale.

**Fig. S3. Pressure effects on phosphatases.** Activity of select phosphatases as indicated without and with 190 kPa added pressure assayed colorometrically from the hydrolysis of p-nitrophenyl phosphate (See Methods).

**Fig. S4. Schematic for study of pressure-induced activity of full length WNK3 in *D. melanogaster* Malphigian tubules.** Endogenous *Drosophila WNK* was knocked down, and full-length human *WNK3* expressed, together with its substrate rat SPAK. Dissected Malphigian tubules were centrifuged to apply hydrostatic pressure, and WNK activity measured by assessing phosphorylation of SPAK using antibodies to phosphorylated (pSPAK) and and total SPAK (tSPAK).

**Fig. S5. Sequence alignments and crosslinking. (A)** Sequence alignment of WNK3, and WNK1 with WNK with selected species, a deep-sea fish (*N. Coriiceps*), shark (*R. Typus*), and a turtle (*P. sinensis*) showing boxes of sequence variation (red) in the dimer interface of uWNK1A. (**B**) Selected crosslinks interpreted as arising from the dimer (black dashed lines) and from the monomer (yellow dashed lines).

**Fig. S6 Cavities in a control kinase** Cavities in (**A**) 3FPQ and (**B**) PAK4 (PDB file 2C30). (**B**) calculated in PYMOL (**B**) The catalytic loop (yellow) and activation loop (red) in PAK4. No water molecules are present between or under these loops, unlike those found in WNK1-KDm (Fig. 4 C,D).

#### Extended Data Table S1. WNK3 autophosphorylation sites

| WNK3 phosphpeptide | Location | 1) From *E. coli* | 2) Dephos | 3) Reautophos |
|---|---|---|---|---|
| $^{1'}$MKET*AAAKF$^{9'}$ | S-Tag | **50%** | **0%** | **30%** |
| $^{10'}$ERQHMDSPDLGTLVPRGS*M$^{28'}$ | Thrombin linker | **90%** | **0%** | **40%** |
| $^{118}$KAVATSPS*GRFL$^{129}$ | Beta sheet 1 | 20% | 20% | 20% |
| $^{283}$ITGPTGS*VKIGDLGLATL$^{300}$ | Beta sheet 7 | 5% | 5% | 10% |
| $^{301}$MRTS*F$^{305}$ | Minor act. loop[1] | **95%** | **30%** | **50%** |
| $^{306}$AKS*VIGTPEFMAPEMY$^{321}$ | Main act. loop[2] | **95%** | **0%** | **80%** |
| $^{359}$RKVTS*GIKPASF$^{370}$ | Helix G | 7% | 0% | 1% |
| $^{404}$FAEDTKLPT*TENLY$^{418'}$ | TEV site | **20%** | **0%** | **2%** |

The phosphorylation state of WNK3-KDm 1) as expressed in *E. coli*, 2) after treatment with PP1Cγ and lambda phosphatase, and 3) after 15 minute *in vitro* phosphorylation reactions of protein from 2). WNK3 protein was proteolysed using chymotrypsin, separated by HPLC and peptides analyzed by mass spectroscopy as described in Supplemental Material. Amino acids in red with "*" denotes phosphorylation sites. AA numbering with ʹ denotes nonphysiological tag sequence (underlined) Numbers indicate percent phosphorylation at each site, with data in blue including canonical activation loop phosphorylation, and green including nonphysiological phosphorylation. [1]Phosphorylation at S304 is the secondary WNK3 activating residue, while [2]Phosphosphorylation at S308 is the canonical primary WNK3 activating residue (reference).

#### Extended Data Table S2. Crosslinks of Unphosphorylated WNK3 mapped onto the WNK1-KDm dimer (6CN9)

| No Pressure Unique X-links dimer | | | | | | |
|---|---|---|---|---|---|---|
| resn 1 | resn 2 | no pr | w pr | Monomer (Å) | Dimer (Å) | |
| 351 | 164 | 18 | n.p. | 31? | 26? | Cat. Loop to N-term |
| 292 | 233 | 15 | n.p. | 18 | 25 | β3 to inside barrel |
| 266 | 164 | 13 | n.p. | 28? | 19? | Hellx C to N-term |
| 381 | 266 | 6 | n.p. | 25 | 19 | Act. Loop to Helix C |

| | | | | | | |
|---|---|---|---|---|---|---|
| 381 | 158 | 10 | n.p. | 37? | 15? | Act loop to N-term |
| No Pressure Unique X-links monomer | | | | | | |
| 322 | 237 | 10 | n.p. | 33 | 53 | β3 to Helix E |
| 233 | 381 | 4 | np | 18 | 43 | 3 to Act. Loop |
| 484 | 333 | 9 | n.p. | 13 | 78 | C-term to Helix E |
| No Pressure Unique X-links, no interpretation | | | | | | |
| 222 | 333 | 9 | n.p. | 38 | 54 | β1 to Helix E |
| Pressure Unique X-links monomer | | | | | | |
| 266 | 233 | n.p. | 13 | 16 | 38 | Helix C to β3 |
| 233 | 317 | n.p. | 8 | 24 | 52 | β3 to DE |
| 256 | 164 | n.p. | 7 | 28? | 14? | N-term to Helix B |
| 310 | 333 | n.p. | 7 | 20 | 56 | Helix D to Helix E |
| 259 | 266 | n.p. | 7 | 11 | 33 | BC to Helix C |
| 434 | 333 | n.p. | 6 | 30 | 51 | Helix E to Helix G |
| 381 | 434 | n.p. | 6 | 18 | 43 | Act. Loop to Helix G |
| 365 | 237 | n.p. | 4 | 19 | 41 | Helix C toβ3 |
| 310 | 222 | n.p. | 4 | 25 | 31 | β1 to Helix D |
| 310 | 233 | n.p. | 3 | 15 | 40 | Helix D to β3 |
| 434 | 237 | n.p. | 11 | 40 | 31 | helix G to β3 |
| Pressure Unique X-links no interpretation | | | | | | |
| 295 | 333 | n.p. | 6 | 36 | 42 | Inside barrel to helix E |

Calculated distances between residue pairs (Resn1-Resn2) in the iWNK1SA structure. Blue indicates distances typical for lysine-lysine DSS crosslinking. Pink indicates distances outside the DSS crosslinking limits. '?' indicates distance to the nearest crystallographically observed residue.

**Extended Data Table S3 Complete list of crosslinks in (A) for unpressurized WNK3 and (B) for pressurized WNK3.**

| Cross linked peptides | Peptide A Residue number | | Peptide B Residue number | | ld-Score |
|---|---|---|---|---|---|
| | WNK3 | WNK1 | WNK3 | WNK1 | |
| FAEDTKLPTTENLY-AEDTKLPTTENLY-a6-b5 | 410 | 484 | 410 | 484 | 22.35 |
| AEDTKLPTTENLY-KGLDTETW-a5-b1 | 410 | 484 | 163 | 237 | 21.13 |
| AKSVIGTPEFMAPEXY-AEDTKLPTTENLY-a2-b5 | 307 | 381 | 410 | 484 | 20.1 |
| LHTRTPPIIHRDLKCDNIF-MKETAAAKF-a14-b8 | 277 | 351 | 90 | 164 | 17.51 |
| AEDTKLPTTENLY-AKSVIGTPEF-a5-b2 | 410 | 484 | 307 | 381 | 17.06 |
| FAEDTKLPTTENLY-FAEDTKLPTTENLY-a6-b6 | 410 | 484 | 410 | 484 | 15.53 |
| DSWESILKGKKCIVLVTELXTSGTLKTY-KTVYKGLDTETWVEVAW-a8-b1 | 218 | 292 | 159 | 233 | 14.73 |
| AKSVIGTPEF-KGLDTETW-a2-b1 | 307 | 381 | 163 | 237 | 14.63 |
| MKETAAAKF-KGLDTETW-a2-b1 | 84 | 158 | 163 | 237 | 13.94 |
| KTVYKGLDTETWVEVAW-MKETAAAKF-a1-b8 | 159 | 233 | 90 | 164 | 13.29 |
| KEEAEXLKGLQHPNIVRFYDSW-XKETAAAKF-a1-b8 | 192 | 266 | 90 | 164 | 12.77 |
| AEDTKLPTTENLYF-AEDTKLPTTENLY-a5-b5 | 410 | 484 | 410 | 484 | 12.51 |
| KGLDTETW-KGLDTETW-a1-b1 | 163 | 237 | 163 | 237 | 12.41 |
| RKVTSGIKPASF-RKVTSGIKPASF-a2-b2 | 360 | 434 | 360 | 434 | 11.72 |
| KVMKPKVLRSW-XKETAAAKF-a1-b2 | 243 | 317 | 84 | 158 | 11.45 |
| KVXKPKVLRSW-KGLDTETW-a6-b1 | 248 | 322 | 163 | 237 | 10.4 |
| RKVTSGIKPASF-MKETAAAKF-a2-b2 | 360 | 434 | 84 | 158 | 10.18 |
| AKSVIGTPEFMAPEXY-MKETAAAKF-a2-b2 | 307 | 381 | 84 | 158 | 9.74 |
| AEDTKLPTTENLYF-CRQILKGLQF-a5-b6 | 410 | 484 | 259 | 333 | 9.34 |
| AKSVIGTPEFXAPEMY-LKFDIELGRGAFKTVY-a2-b2 | 307 | 381 | 148 | 222 | 9.22 |
| DIELGRGAFKTVY-XKETAAAKF-a10-b2 | 159 | 233 | 84 | 158 | 9.07 |
| LKFDIELGRGAF-CRQILKGLQF-a2-b6 | 148 | 222 | 259 | 333 | 8.7 |
| LKRFKVXKPKVLRSW-AEDTKLPTTENLYF-a5-b5 | 243 | 317 | 410 | 484 | 7.49 |
| DIELGRGAFKTVY-MKETAAAKF-a10-b8 | 159 | 233 | 90 | 164 | 7.38 |
| KTVYKGLDTETW-XKETAAAKF-a5-b8 | 163 | 237 | 90 | 164 | 6.87 |
| AKSVIGTPEFMAPEXYEEHY-KEEAEXLKGLQHPNIVRF-a2-b1 | 307 | 381 | 192 | 266 | 6.54 |
| AKSVIGTPEFMAPEXY-AKSVIGTPEFXAPEMY-a2-b2 | 307 | 381 | 307 | 381 | 5.93 |
| AKSVIGTPEFMAPEXYEEHY-LKFDIELGRGAFKTVY-a2-b13 | 307 | 381 | 159 | 233 | 4.07 |
| AEDTKLPTTENLY-AEDTKLPTTENLY-a5-b5 | 410 | 484 | 410 | 484 | 3.38 |

Extended Data Table 3B

B

| Cross linked peptides | Peptide A Residue Number | | Peptide B Residue Number | | ld-Score |
|---|---|---|---|---|---|
| | WNK3 | WNK1 | WNK3 | WNK1 | |
| KGLDTETW-KGLDTETW-a1-b1 | 163 | 237 | 163 | 237 | 29.47 |
| FAEDTKLPTTENLY-FAEDTKLPTTENLY-a6-b6 | 410 | 484 | 410 | 484 | 20.97 |
| AEDTKLPTTENLY-AKSVIGTPEF-a5-b2 | 410 | 484 | 307 | 381 | 20.27 |
| FAEDTKLPTTENLY-AEDTKLPTTENLY-a6-b5 | 410 | 484 | 410 | 484 | 18.43 |
| AKSVIGTPEFMAPEXY-AEDTKLPTTENLY-a2-b5 | 307 | 381 | 410 | 484 | 18.16 |
| AKSVIGTPEFXAPEMY-AEDTKLPTTENLY-a2-b5 | 307 | 381 | 410 | 484 | 18.02 |
| AEDTKLPTTENLY-KGLDTETW-a5-b1 | 410 | 484 | 163 | 237 | 16.44 |
| AEDTKLPTTENLY-AEDTKLPTTENLY-a5-b5 | 410 | 484 | 410 | 484 | 16.2 |
| XKETAAAKF-KGLDTETW-a2-b1 | 84 | 158 | 163 | 237 | 15.86 |
| LKFDIELGRGAF-XKETAAAKF-a2-b2 | 148 | 222 | 84 | 158 | 15.53 |
| DIELGRGAFKTVY-MKETAAAKF-a10-b8 | 159 | 233 | 90 | 164 | 15.26 |
| KGLDTETWVEVAW-AKSVIGTPEF-a1-b2 | 163 | 237 | 307 | 381 | 14.78 |
| AKSVIGTPEF-XKETAAAKF-a2-b8 | 307 | 381 | 90 | 164 | 14.67 |
| KEEAEXLKGLQHPNIVRFY-DIELGRGAFKTVY-a1-b10 | 192 | 266 | 159 | 233 | 13.27 |
| XKETAAAKF-XKETAAAKF-a2-b8 | 84 | 158 | 90 | 164 | 12.39 |
| KEKNEKEMEEEAEXKAVATSPSGRF-CRQILKGLQF-a1-b6 | 122 | 196 | 259 | 333 | 11.05 |
| KTVYKGLDTETW-RKVTSGIKPASF-a5-b2 | 163 | 237 | 360 | 434 | 10.57 |
| LKFDIELGRGAFKTVY-MKETAAAKF-a2-b8 | 148 | 222 | 90 | 164 | 10.54 |
| RKVTSGIKPASF-MKETAAAKF-a2-b2 | 360 | 434 | 84 | 158 | 10.53 |
| KGLDTETWVEVAW-KGLDTETW-a1-b1 | 163 | 237 | 163 | 237 | 10.12 |
| AEDTKLPTTENLYF-AEDTKLPTTENLYF-a5-b5 | 410 | 484 | 410 | 484 | 10.07 |
| AEDTKLPTTENLY-LKFDIELGRGAF-a5-b2 | 410 | 484 | 148 | 222 | 8.46 |
| DIELGRGAFKTVY-KVXKPKVLRSW-a10-b1 | 159 | 233 | 243 | 317 | 8.46 |
| CELQDRKLTKAEQQRF-MKETAAAKF-a7-b8 | 182 | 256 | 90 | 164 | 7.38 |
| DSWESILKGKKCIVLVTELXTSGTLKTY-CRQILKGLQF-a26-b6 | 236 | 310 | 259 | 333 | 6.73 |
| CELQDRKLTKAEQQRFKEEAEXLKGLQHPNIVRF-KEEAEMLKGLQHPNIVRFYDSW-a10-b1 | 185 | 259 | 192 | 266 | 6.58 |
| LKFDIELGRGAFKTVY-MKETAAAKF-a13-b2 | 159 | 233 | 84 | 158 | 6.58 |
| SECQNAAQIYRKVTSGIKPASF-CRQILKGLQF-a12-b6 | 360 | 434 | 259 | 333 | 5.94 |
| ESILKGKKCIVLVTELMTSGTLKTYLKRF-CRQILKGLQF-a8-b6 | 221 | 295 | 259 | 333 | 5.67 |
| AKSVIGTPEFMAPEMY-RKVTSGIKPASF-a2-b2 | 307 | 381 | 360 | 434 | 5.63 |
| AKSVIGTPEFMAPEXYEEHY-LKFDIELGRGAFKTVY-a2-b2 | 307 | 381 | 148 | 222 | 4.87 |
| ITGPTGSVKIGDLGLATLMRTSF-KTVYKGLDTETWVEVAW-a9-b5 | 291 | 365 | 163 | 237 | 4.4 |
| FAEDTKLPTTENLY-AKSVIGTPEF-a6-b2 | 410 | 484 | 307 | 381 | 3.9 |
| ESILKGKKCIVLVTELMTSGTLKTY-LKFDIELGRGAF-a23-b2 | 236 | 310 | 148 | 222 | 3.89 |
| ESILKGKKCIVLVTELMTSGTLKTY-DIELGRGAFKTVY-a23-b10 | 236 | 310 | 159 | 233 | 2.66 |
